# The weighting is the hardest part: on the behavior of the likelihood ratio test and score test under weight misspecification in rare variant association studies

**DOI:** 10.1101/020198

**Authors:** Camelia C. Minică, Giulio Genovese, Christina M. Hultman, René Pool, Jacqueline M. Vink, Conor V. Dolan, Benjamin M. Neale

## Abstract

Rare variant association studies are at a critical inflexion point with the increasing availability of exome-sequencing data. A popular test of association is the sequence kernel association test (SKAT). Weights are embedded within SKAT to reflect the hypothesized contribution of the variants to the trait variance. Correct weighting is expected to boost power, and yet the correct weights are generally unknown. It is therefore important to assess the effect of weight misspecification in SKAT.

We evaluated the behavior of the score and likelihood ratio tests (LRT) under weight misspecification. Simulation and empirical results revealed that LRT is generally more robust and more powerful than score test in such a circumstance. For instance, when the simulated betas were larger for rarer than for more common variants, (incorrectly) assigning equal weights reduced the power of the LRT by ∼ 5%, while the score test’s power dropped by ∼ 30%.

To optimize weighting we proposed a data-driven weighting scheme. With this scheme and LRT we detected significant enrichment of rare case mutations (MAF<5%; P-value=7e-04) of a set of constrained genes in the Swedish schizophrenia case-control cohort with exome-sequencing data.

The score test is currently preferred for its computational efficiency and power. Indeed, assuming correct specification, in some circumstances the score test is the most powerful test. However, LRT has the compelling qualities of being generally more powerful and more robust to misspecification. This is an important result given that, arguably, misspecified models are likely to be the rule rather than the exception in weighting-based approaches.

## 1. Introduction

With the availability of high-coverage exome/genome sequence data in increasingly large samples, rare variant association studies (RVAS) are gaining importance in human genetic research. One important test of association between a target set of rare variants (RVs) and a given phenotype is the sequence kernel association test (SKAT; [1, 2, 3, 4, 5, 6, 7]). SKAT is based on a random effects model, in which the effect sizes of the RVs are assumed to be drawn from a zero mean distribution and variance that can be specified by weights. These weights are typically assigned based on meta-information about the RVs, such as allele frequency and functional predictions [8, 9, 10, 1], with rarer and functional variants expected to have larger effects. Allele frequency, in particular, is an important weighting factor, as the rarer the variant is, the stronger the average purifying selection coefficient [11, 12]. If this assumption is true, the effect sizes for rare variants will tend to be larger than for more common variants.

The relationship between effect size, frequency and selection, however, rests on assumptions about the extent of direct selection on the phenotype in question and the demographic history of the population [13, 10, 14]. Genomic regions under low selection pressures may harbor rare as well as more common so-called goldilocks alleles, both with strong functional effects, as simulation studies [10] and empirical results have demonstrated (e.g., [15]). Testing such genomic regions by relying on a weighting scheme which up-weighs rarer variants and puts low or zero weights on the more common ones, may weaken the association signal. Correct weighting is expected to boost the power of detection [1]. However, as the true weights are generally unknown, it is important to establish the effect of weight misspecification in a kernel-based variance component test. Here we assessed the loss of power associated with incorrect weighting in sequence-based kernel association tests. Because hypothesis testing can be performed by using either the score [1] or the likelihood ratio test [5], we characterize the behavior of both tests within the misspecification space. We considered various weighting schemes and target regions harboring functional variants or mixtures of functional and neutral variants. We show that the choice of the statistical test has an important bearing on power, with the likelihood ratio test being appreciably more robust to weight misspecification. Furthermore, we show that the power loss depends not only on the degree of misspecification, but also on the presence of neutral variants within the target set. As to how to minimize the power loss resulting from misspecification of weights, we examined the efficiency of a data-driven weighting scheme. We propose the use of a set of theoretically defensible weighting schemes, of which, we assume, the one that gives the largest test statistic is likely to capture best the allele frequency-functional effect relationship. The use of alternative weighting schemes is intended to accommodate genomic regions where only very rare variants are likely to be functional, as well as regions under weak selection pressures, harboring both rare and common variants, both (possibly) related to the risk of the disease of interest. Family-wise error rate can be protected either by permutations or by using a Bonferroni correction. We show the power benefits conferred by the use of such a variable data-driven weighting procedure both in simulated and in empirical data.

Below we first formulate the model and briefly consider the two tests of variance components, namely the likelihood ratio test and the score test. Next we explore the behavior of the two tests under (in)correct model specification in a simulation study. We then present and evaluate the use of a data-driven weighting scheme in simulated and empirical data. Finally, we discuss the robustness of the likelihood ratio test to misspecification and the power advantages conferred by our proposed weighting procedure in SKAT.

## 2. Materials and Methods

### 2.1. Model formulation

Let **y** be the *n*-dimensional vector of continuous phenotypes measured in a sample consisting of *n* individuals. Let **X** be the *n × p* design matrix containing the relevant covariates. Let **G** be the *n × m* matrix of genotype values, with the *g*_*ij*_ element denoting the genotype value of the individual *i* (*i* = 1 *… n*) at locus *j* (*j* = 1 *… m*). Genotypes are coded as additive-codominant, i.e., *g*_*ij*_ = (0, 1, 2). The association between the phenotype and the set of *m* SNPs is modeled within the linear mixed model framework as:

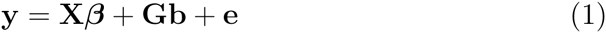

with ***β***^*t*^ = (*β*_1_, *… β_p_*) being the *p*-dimensional vector of fixed effects of co-variates, **b**^*t*^ = (*b*_1_, *…, b_m_*) being the *m ×* 1 vector of regression coefficients in the regression of the phenotype on the *m* genetic variants within the target set, and **e** being the *n*-dimensional vector of random residuals. The random *b* vectors **b** and **e** are assumed to be normally distributed: 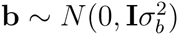 and 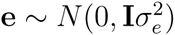, with **I** being the identity matrix of appropriate dimension.

Let **W** be the *m × m* diagonal matrix containing the weights used to weigh the contribution to the test statistic of the SNPs in the set. The normally distributed phenotype **y** has expected mean **E**[**y**] = **X*β*** and variance-covariance matrix:

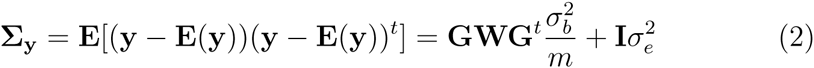

with **GWG**^*t*^ being the weighted kernel or genetic relationship matrix. As implemented in the SKAT [1], the diagonal elements of the matrix **W**, diag(*w*_1_ *…, w_m_*), are related to the minor allele frequency of the *j*-th variant by means of the beta density distribution function (dbeta), which is characterized by two shape parameters. The specification of the two shape parameters is informed by the hypothesized relationship between the *j*-th variant effect and its minor allele frequency (MAF; see section on ‘Weighting’ below).

### 2.2. Tests of variance components

To test whether the parameter of interest 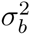 deviates significantly from zero, one can employ a likelihood ratio test (LRT) or a score test. The likelihood ratio test is computed as two times the difference between the log-likelihoods of the null model (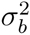 constrained to equal 0) and the alternative model (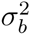 estimated freely). Parameter estimation can be performed by restricted/residual maximum likelihood (REML):

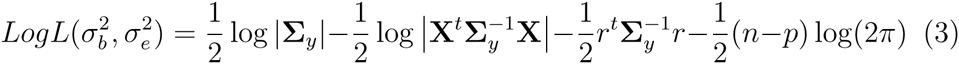

where 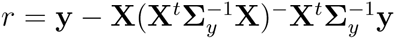 with superscript ‘–’ denoting a generalized inverse [16].

In evaluating the statistical significance of the restricted LRT, we note that the null distribution of the test statistic is an equally weighted. 5 :. 5 mixture of a 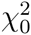 and a 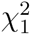 distributions (see e.g., [17, 18, 19]). Alternatively, the null distribution can be constructed empirically by using a permutation-based approach (e.g., [5]), or a parametric bootstrap (e.g., [20]).

The score test is computed as:

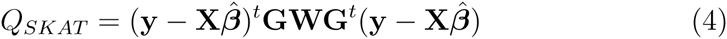

with its expected null distribution following a mixture of chi-square distribution and statistical significance assessed by means of the Davies exact method [21].

### 2.3. Data simulation

Phenotypes and genotypes were generated in samples of *n* = 10, 000 un-related individuals. Specifically, we simulated two *m*-dimensional random vectors of continuous variables representing alleles at *m* equidistant loci for each individual *i* from the sample. The vectors were drawn from a multi-variate distribution with zero mean and **Σ**_*LD*_ correlation matrix. The SNPs were in linkage equilibrium, i.e., we set **Σ**_*LD*_ to equal an identity matrix. The multivariate normally distributed variables were then discretized given chosen thresholds based on the MAF at each locus. We considered MAFs varying randomly between 0.005 and 0.05, sampled from a uniform distribution. Given the vectors of alleles, we then created the *m* vectors of genotypes, *g*_*ij*_. Based on the genotypes, the *n* × 1 vector of phenotypes, **y**, was generated as:

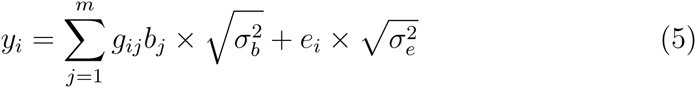

*b*_*j*_, the regression weight of the SNP at the *j*-th locus, was computed as a function of MAF_*j*_ and of its contribution to the standardized variance of the polygenic scores [22]. Namely, the regression weights varied with MAF, while their contribution to the genetic variance was equal. Simulating data in this fashion is equivalent to simulation according to dbeta(MAF,. 5,.5) weights [1], with weights increasing with decreasing MAF. We also simulated data according to dbeta (1,1) weights (second simulation scenario), where SNPs had equal weights regardless of MAF. This scenario is illustrative for situations where the tested region harbors both common and rare variants, both having functional effects on the trait (i.e., where there is no relationship between allele frequency and effect size). The variance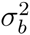 equaled 0.01 across *eb* all scenarios we considered, and 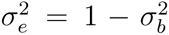. The *n*-dimensional vector of environmental scores **e** was drawn from a standard normal distribution *N* (0, 1).

### 2.4. Exploring the misspecification space: Weighting

To explore the effect of weight misspecification on the power and type I error rates of the LRT and the score test we carried out simulations. The *m*-dimensional vector **w** of SNP weights was computed using the beta density function, with the *j*-th element calculated as *w*_*j*_ = *dbeta*(*M AF_j_*; *a*_1_, *a*_2_) given the MAF of the *j*-th variant and the shape parameters *a*_1_ and *a*_2_. As described in the previous section, data were simulated according to: a) dbeta(.5,.5) weights (i.e., the true weights increase with decreasing MAF); and b) dbeta(1,1) weights (i.e., the SNPs have equal weights, regardless of MAF). Next, in computing the tests statistic we (mis)specified the weights as: a) dbeta(1,1); b) dbeta(.5,.5); c) dbeta(1,25). The first weighting scheme pertains to the hypothesis that there is no relationship between the regression weight and the frequency of the variant (hence, the more common variants contribute on average more to variation in the phenotype). In this scenario the association test is carried out with raw additive-codominant coding of the genotypes. The use of the second weighting scheme is equivalent to standardization of the genotypic values prior to the analysis. We considered the effect of this weighting scheme as this treatment of the genotypes is default in GCTA [23] and in FaST-LMM-set [5]. Standardization and assignment of weights dbeta(.5,.5) are equivalent weighting schemes [1] in which the contribution to the test of rarer variants is up-weighed relative to that of the more common ones [24], and hence the variants contribute on average equally to the variance in the phenotype, regardless of frequency. We also considered the effects of the third weighting scheme (dbeta(1,25)) as weights computed as such are the default weights in SKAT [1].

We assessed the behavior of the two tests under weight misspecification by considering: a) target regions harboring solely functional variants with opposite effects on the phenotypic mean, and b) regions harboring a mixture of protective, deleterious and neutral effects.

### 2.5. Evaluating the type I error rates and power

We evaluated the type I error rates by generating 1,000,000 datasets under the null hypothesis of no phenotypic variance explained by the SNPs within the target set. The type I error rate was computed as the proportion of datasets in which the tests incorrectly rejected the null hypothesis and it was evaluated given *α*=0.01 and 0.001.

Power was assessed based on 1000 simulated datasets, an effect size of 1% explained phenotypic variance and 7 alpha thresholds. Given the 7 alpha thresholds, power equaled the proportion of datasets in which the effect was detected. As a validity check of our program, for all the scenarios considered we also report the power and the type I error rates of the true (i.e., correct) model.

### 2.6. Variant weighting schemes: data-driven search for optimal weights

Because the application of a single weighting scheme might not be accurate when testing thousands of genes scattered across the whole exome (possibly subjected to selection pressures of varying intensities) we also considered the efficiency of a data-driven search for the optimal weights. We generated 1000 samples according to weights dbeta(.5,.5) as described above. Each generated sample comprised N=3000 individuals with phenotypes and genotypes observed at 50 variant sites. As above, the variants were either all functional (deleterious or protective, first scenario) or a mixture of functional and neutrals (second scenario). We performed association tests by using a set of 7 weighting schemes: a) dbeta(1,75) ; b) dbeta(1,50); c) dbeta(1,35); d) dbeta(1,25); e) dbeta(1,5); f) dbeta(1,1) and g) dbeta(.5,.5). Statistical significance was assessed by means of permutations on phenotypes. Specifically, we computed a maximum test statistic max_*LRT*_ (max_score_) as the largest out of the seven tests obtained given the genotypes transformed according to each of the weighting schemes enumerated earlier. We then repeated this step in 1000 permuted datasets obtained by shuffling the phenotypes. We computed the p-value as the proportion of datasets in which the max_*LRT*_ (max_score_) was larger in the permuted than in unpermuted data. Power equalled the proportion of datasets yielding a p-value smaller than 0.01.

### 2.7. Software

The R-package MASS [25] was used for data generation. Model fitting was performed in R-nlme [26], and SKAT [27]. We used the anova function in R to obtain the restricted likelihood ratio test, with the p-value computed by halving the supplied p-value [28]. To check our model fitting approach, we analyzed one simulated sample of 10,000 individuals by using 3 independent programs implementing genetic similarity/kernel-based variance component tests: the nlme R-package, the software Genome-wide Complex Trait Analysis (GCTA; [23]) and the software FaST-LMM-set [5]. The values for the restricted LRT and the estimate for the variance component obtained by the 3 programs were almost identical (see Table 1 Supplementary Material for details https://goo.gl/Thz2cM), indicating that these are equivalent approaches. Having established the equivalence, all the simulations were next conducted using the nlme program. Simulations were carried out on the Broad Institute Gold Compute cluster.

**Table 1:** Type I error for the restricted likelihood ratio test (LRT) and the score test, given genotypic data simulated under the null model of no association between the target region and the phenotype. The sample consisted of 10,000 individuals with genotypes at 50 SNPs having minor allele frequencies (MAFs) sampled from the uniform distribution and ranging from. 5% to 5%. The restricted LRT and the score tests were computed for three sets of weights beta in each of the 1,000,000 simulated samples. Type I error equals the proportion of datasets in which the null hypothesis has been incorrectly rejected given the three significance thresholds.

| | weights<br>dbeta | $\alpha=0.01$ [99%CI] | $\alpha=0.001$ [99%CI] |
| --- | --- | --- | --- |
| **LRT** | (.5,.5) | 0.00849 [0.00826, 0.00873] | 0.00083 [0.00076, 0.00091] |
|  | (1,1) | 0.00834 [0.00811, 0.00858] | 0.0008 [0.00073, 0.00088] |
|  | (1,25) | 0.00847 [0.00824, 0.00871] | 0.00082 [0.00075, 0.0009] |
| **Score** | (.5,.5) | 0.00991 [0.00966, 0.01017] | 0.00098 [0.0009, 0.00107] |
|  | (1,1) | 0.00992 [0.00967, 0.01018] | 0.00099 [0.00091, 0.00107] |
|  | (1,25) | 0.00988 [0.00962, 0.01013] | 0.00098 [0.0009, 0.00106] |

**Table 2:** Type I error for the restricted likelihood ratio test (LRT) and the score test, given genotypic data simulated under the null model of no association between the target region and the phenotype. The sample consisted of 10,000 individuals with genotypes at 50 SNPs having equal beta weights and minor allele frequencies (MAFs) sampled from the uniform distribution and ranging from. 5% to 5%. The LRT and the score tests were computed for three sets of weights beta in each of the 1,000,000 simulated samples. Type I error equals the percent of datasets in which the null hypothesis has been incorrectly rejected given the three significance thresholds.

| | weights<br>dbeta | $\alpha=0.01$ [99%CI] | $\alpha=0.001$ [99%CI] |
| --- | --- | --- | --- |
| **LRT** | (.5,.5) | 0.00844 [0.00821, 0.00868] | 0.0008 [0.00073, 0.00088] |
|  | (1,1) | 0.00844 [0.00821, 0.00868] | 0.0008 [0.00073, 0.00088] |
|  | (1,25) | 0.00817 [0.00794, 0.00841] | 0.00074 [0.00067, 0.00082] |
| **Score** | (.5,.5) | 0.00989 [0.00964, 0.01015] | 0.00099 [0.00091, 0.00107] |
|  | (1,1) | 0.0098 [0.00954, 0.01005] | 0.00098 [0.00090, 0.00106] |
|  | (1,25) | 0.00993 [0.00968, 0.01019] | 0.00094 [0.00086, 0.00102] |

### 2.8. Empirical analysis: testing the constrained and the FMRP-Darnell gene sets for rare case mutations enrichment

We compared the performance of the likelihood ratio test and of the score test under alternative weights in a real dataset. For this illustration we used the Swedish schizophrenia case-control cohort of 4940 individuals with exome-sequencing data from blood DNA. Cases had a clinical diagnosis of schizophrenia and at least two hospitalizations as determined by expert review based on the Hospital Discharge Register [29, 30]. Controls, without a diagnosis of schizophrenia or bipolar disorder, were randomly selected from population registries. Both cases and controls are of Scandinavian ancestry, aged 18 or older (see [31, 32] for a detailed description of the sample). There were 169 individuals with unreliable samples (i.e., duplicates, ethnic outliers or having a genotype missing rate higher than 10%) whom we removed from the analysis. This left for the analysis 2461 cases and 2479 controls. 2732 of these were males. Written informed consent was obtained from all participants (or legal guardian consent and subject assent). All procedures were approved by the ethical committees in Sweden and in the United States.

Exome-sequencing was performed in seven waves at the Broad Institute of MIT and Harvard. For samples in the first wave, hybrid capture was performed using the Agilent SureSelect Human All Exon Kit method. In this version, the method targets *∼* 28 million base-pairs partitioned in *∼* 160,000 regions. Sequencing was done using Illumina GAII instruments. For samples in the waves two to seven, hybrid capture was done by using the newer version of the Agilent SureSelect Human All Exon v.2 Kit method, which targets ~32 million base-pairs partitioned in *∼* 190,000 regions. Sequencing was performed using the Illumina HiSeq 2000 and HiSeq 2500 instruments. We used BWA ALN version 0.5.9 [33] to align the reads to the GRCh37 human genome reference and we applied Picard/GATK to process the sequence data and to call variants http://broadinstitute.github.io/picard/; [34]. Selected singletons were validated using Sanger sequencing (see [32] for details).

Variants out of Hardy-Weinberg equilibrium (P-value *<* 5e-8) and showing excess heterozygosity, or variants showing excessive correlation (P-value *<* 5e-8) with the covariates (that could not be explained by principal components) were excluded from the analysis. In addition, we excluded variants that did not pass the GATK default filters [35, 36]. There were 892,306 variants with MAF *<* 5% meeting all our quality control criteria.

For this empirical illustration we considered the gene-sets rather than the genes as the unit of analysis. The reason that we extended the targeted region is that the current sample sizes afford insufficient power for gene-based tests (see Purcell et al., 2014) but are more adequate for gene-set enrichment analyses which consider jointly a larger number of weak effects. This type of analysis has the added benefit of reducing substantially the burden of multiple testing. By extending the targeted region, the number of tested variants is large, and hence the effects of (possible) weight misspecification are expected to be large. In addition, as we do not focus on a specific class of alleles but rather lump together all observed variants with frequency below specific thresholds, a large amount of variation contributing to the test statistic will possibly be neutral. This makes the example a near optimal situation for illustrating the difference in robustness to both model misspecification and neutral variation of the LRT and the score test.

We tested for enrichment of case mutations two partially overlapping gene-sets likely relevant to schizophrenia. The first set consisted of 899 genes which are part of the list identified by Samocha et al. [37] as highly constrained. These constrained genes were proposed as candidates in autism spectrum disorder (ASD) given their enrichment for de novo loss of function case mutations. Given evidence favouring the hypothesis that schizophrenia and ASD share genetic aetiology [38, 39], this set of genes is likely to be relevant also to schizophrenia. The second set consisted of 749 genes targeted by the Fragile-X mental retardation protein (FMRP). This set is part of the list of genes derived by Darnell et al. [40] from mouse brain as likely implicated in regulating synaptic plasticity. Genes targeted by FMRP were found to be enriched for de novo nonsynonimous case mutations in both ASD [41] and schizophrenia [39]. Purcell et al. (2014) also tested the FMRP set for enrichment of rare variants in the current sample, and their analysis yielded nominally significant results. Note that the strategy we adopted here is however, different. That is, rather than using gene-based statistic, our procedure tests for the joint effect (variance explained) of rare variants with MAF lower than 5% and 1% within the gene-set (note that the MAF thresholds are, however, arbitrary: variants defined as rare in one sample might feature as common in another sample).

We performed sequence-based kernel association analyses using the likelihood ratio and score tests with variable weights. For this empirical analysis we used the FaST-LMM-Set software [5]. To adjust for ancestry we included into analysis the first two principal components. Principal components were computed from genotypes at variants shared with the 1000 Genomes Project phase 1 dataset. To accommodate the scenario in which only very rare variants are likely to be functional, as well as the scenario in which the targeted region is under weak selection pressures, harboring both rare and more common variants, both (possibly) related to the risk of disease (regardless of frequency), we used three alternative weighting schemes: dbeta(1,25), dbeta(.5,.5) and dbeta(1,1). To reduce the computational burden, we chosen to adapt our alpha for multiple testing rather than to rely on permutations to compute the p-value. Hence for each tested pathway, we chose the p-value corresponding to the weighting scheme that yields the largest test statistic. An alpha of 0.05/12= 0.004 was used, corrected for multiple hypothesis testing of 2 gene-sets, 2 frequency thresholds and 3 weighting schemes. For computational ease we used a linear model [5]. The linear LRT (and the linear score test) shows good control of the type I error rate and has performed as well as a generalized linear model in case-control samples (see [6]).

## 3. Results

### 3.1. Type I error

Tables 1 and 2 contain the results pertaining to the type I error rates of the two tests, given correct and incorrect model specification. Across all conditions evaluated here, the score test shows good control of the type I error rate. The likelihood ratio test appears slightly conservative, regardless of whether the weights are correctly specified or misspecified. A similar result was reported by Listgarten et al. [5] who suggested that relying on a. 5:.5 mixture of a 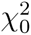 and a 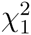 distribution to assess statistical significance of the one variance component LRT might be conservative. We used this approach in the simulations as this is default in most statistical software (e.g., in GCTA, [23]). Alternatively, Listgarten et al. (2013) proposed a permutation based approach to construct the null distribution of the test statistic, approach that maintains the type I error rate of the restricted LRT closer to the expectation. This approach, however, is computationally demanding especially when the number of tested variants within the target and the sample is large.

### 3.2. Power

Figure 1 and Figure 2 display the results relating to power. Four important conclusions follow from our simulation results. First, the restricted LRT and the score test have equal power under correct weight specification. This is expected, as the two tests are asymptotically equivalent when the model is true, i.e., correctly specified (e.g., [42]). The powers of the two tests – displayed in grey in the power figures – are indistinguishable when the assigned weights correspond to the true weights.

**Figure 1:**
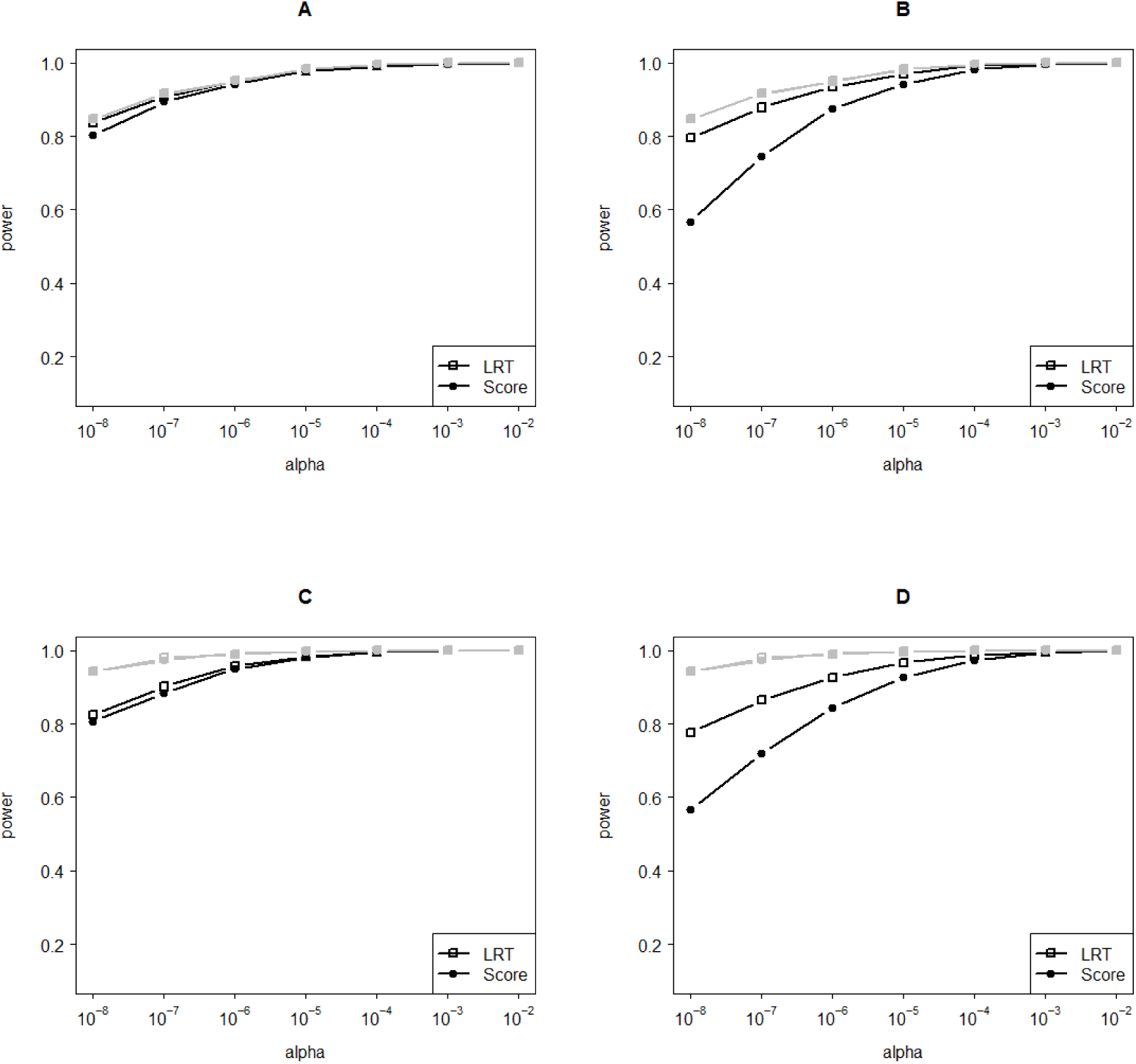
The power of the likelihood ratio test (LRT) and the score test to detect a gene harboring 50 low-frequency SNPs: all functional (A and B) or a mixture of 30 functional and 20 neutral SNPs (C and D). We randomly sampled MAFs ranging from. 5% to 5% from the uniform distribution. The gene explains 1% of the phenotypic variance. Genotypic data were simulated according to weights dbeta(.5,.5), models were fitted according to weights dbeta(1,25) (A and C) and dbeta(1,1) (B and D). The power of the true models (i.e., with correct weights) is displayed in grey. Power was evaluated in 1000 datasets consisting of 4000 individuals. Note that while the SNP-set explain the same amount of phenotypic variance (i.e., 1%) across all scenarios considered, the true individual SNP weights increase as the proportion of functional SNPs in the set decreases.

**Figure 2:**
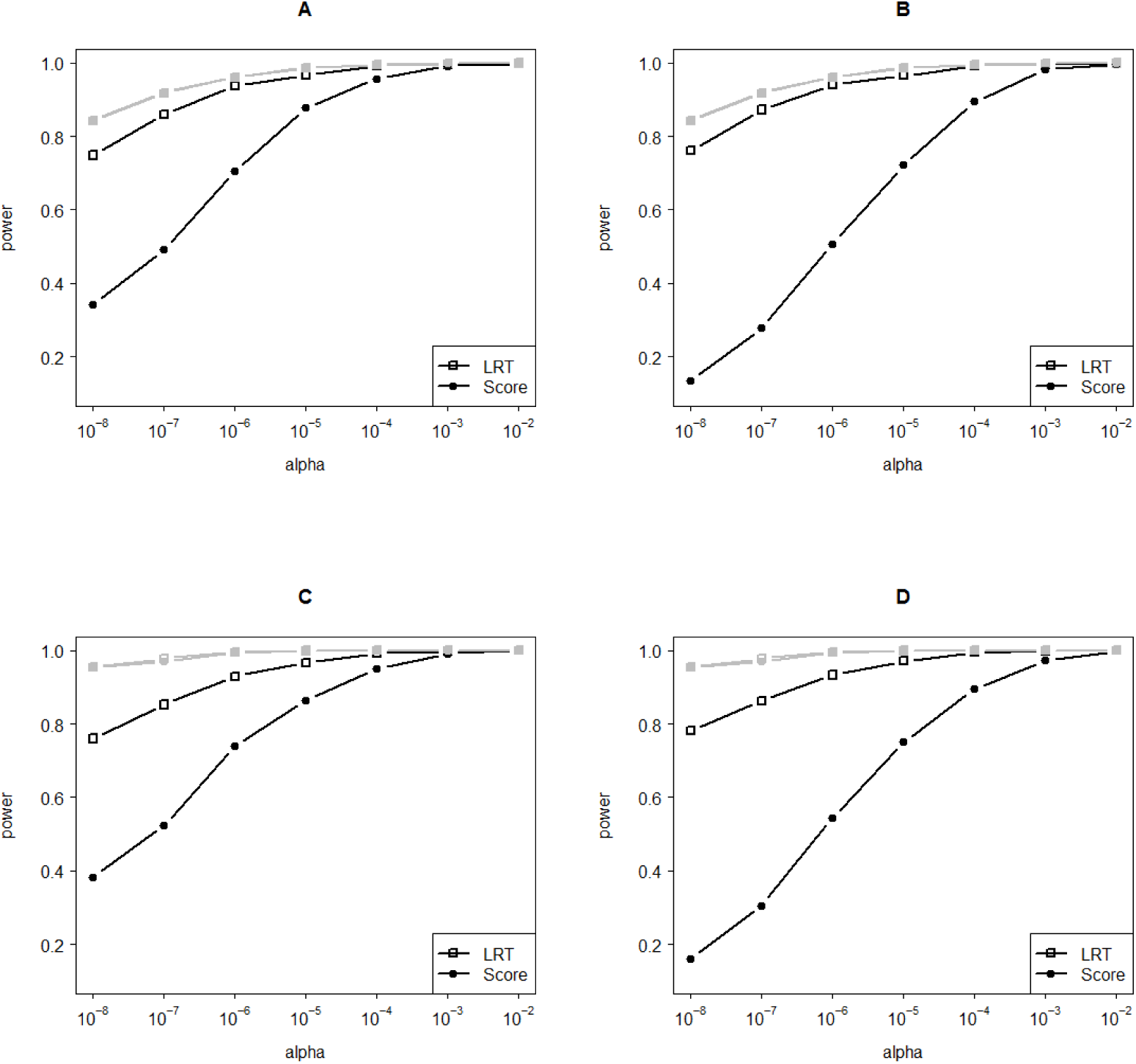
The power of the likelihood ratio test (LRT) and the score test to detect a gene harboring 50 low-frequency SNPs: all functional (A and B) or a mixture of 30 functional and neutral SNPs (C and D). We randomly sampled MAFs ranging from. 5% to 5% from the uniform distribution. The gene explains 1% of the phenotypic variance. Genotypic data were simulated according to weights dbeta(1,1), models were fitted according to weights dbeta(1,25) (A and C) and dbeta(.5,.5) (B and D). The power of the true models (i.e., with correct weights) is displayed in grey. Power was evaluated in 1000 datasets consisting of 4000 individuals. Note that while the SNP-set explain the same amount of phenotypic variance (i.e., 1%) across all scenarios considered, the true individual SNP weights increase as the proportion of functional SNPs in the set decreases.

Second, misspecification of weights always reduces power. This is shown in Figure 1 and in Figure 2, as the departure of the power under model mis-specification (the black lines) from the power of the true models (the grey lines). The exact loss in power depends on the degree of weight misspecification and on the statistical test employed. We note that the power loss is relatively small given mild misspecification of weights. This result is illustrated in Figure 1A, where the assigned weights dbeta(1,25) resemble the true weights dbeta(.5,.5). In this circumstance, it is mainly the presence of neutral SNPs in the target that dilutes the power (see Figure 1C). However, the power may suffer dramatically with increasing misspecification. For instance, when data were simulated according to the dbeta(.5,.5) weights, using a dbeta (1,1) weighting scheme (equal weights assigned to all variants) results in a loss in power of up to *∼*5% and *∼*30% for the restricted LRT and for the score test, respectively (see Figures 1B and 1D). This result is informative for RVASs in which the raw genotypes (unweighted) are used in the test of association. A more dramatic power loss is illustrated in Figure 2D where we consider the reverse situation: weights dbeta (.5,.5) are assigned to SNPs simulated under flat weights. That is, in this scenario, the allele frequency is incorrectly used to inform on the weights assignment. With this misspecification the drop in power relative to the true model is ∼17% and ∼ 80% for the restricted LRT and for the score test, respectively. Third, the inclusion of neutral SNPs dilutes the power of both tests. In our examples, with 40% neutral SNPs the power drops are in the range of ∼10%-∼ 17% relative to the power of the true model, regardless of the degree of weight misspecification. Clearly, discarding neutral variation present within the target is beneficial to improve power to detect significant association.

Forth, relative to the score test, we note that the restricted LRT is consistently more robust, both to weight misspecification and to the presence of neutral variation in the target region. These results are consistent with those reported by Lippert et al. [6], who found their proposed LRT to be generally more powerful than the score test across their simulated settings. Although Lippert et al. did not consider the behavior of the two tests under misspecified weights, they reported the same pattern of results in real data analysis, where the LRT yielded consistently more associations than the score test. As the real weights are in all likelihood not known, the superior power of the restricted LRT in real data might be explained as well by its robustness to weight misspecification and to the inclusion of weighed neutral variation in the computation of the test statistic.

### 3.3. Variable weighting schemes: data-driven search for optimal weights

Table 3 contains the simulation results relating to the power of the two tests under alternative weighting schemes. On one hand, the likelihood ratio test appears to benefit from the use of variable weights. When the variants within the target are all functional, the use of alternative weights increases its power relative to the incorrect weighting setting (power goes up from 48.3% to 52.9%). This increase is to be larger (about 7% given alpha of 0.01) when neutral variants are also present in the target set. On the other hand, the score test performs worst in the variable weighting setting. Power goes down about 10% relative to the incorrect weighting setting, both when the target set contains only functional variants or a mixture of functional and neutral variants.

**Table 3:** Power results for the variable threshold approach given all variants being functional in the target set (A) or a mixture of protective, deleterious and neutral variants (B). Power was evaluated given alpha of 0.01 in 1000 simulated samples, each sample comprinsing 3000 individuals. The minor allele frequency of the 50 variants within the target set ranged from. 5% to 5%.

|  | **weights**<br>**dbeta** | **LRT** | **Score** |
| --- | --- | --- | --- |
| A. | (1,1) (Incorrect) | 0.483 | 0.469 |
|  | Variable weights | 0.529 | 0.364 |
|  | (.5,.5) (True) | 0.547 | 0.559 |
|  | **weights**<br>**dbeta** | **LRT** | **Score** |
| B. | (1,1) (Incorrect) | 0.498 | 0.471 |
|  | Variable weights | 0.564 | 0.382 |
|  | (.5,.5) (True) | 0.709 | 0.734 |

It should be noted, however, that there is a price to pay in terms of power by using a variable weighting scheme in contrast to correct weighting. The price is largest for regions containing mixtures of functional and neutral variants (e.g., the power of the LRT decreases from 70.9% given correct weights to 56.4% with the variable weighting approach) and relatively small for the (less realistic) scenarios in which the target set contains only functional variants (i.e., with the LRT, the power drops about 2%).

As typically the true weights are unknown, conjecturing the correct ones by employing alternative weights and using the likelihood ratio test appears to be the strategy likely to maintain the power close to that of the true model. This strategy appears to be advantageous especially when the target set contains also neutral variants. However, by being based on permutations, the variable weighting approach is likely to be computationally too complex when the number of tests and the sample is large.

### 3.4. Empirical analysis: testing the constrained and the FMRP-Darnell gene sets for rare case mutations enrichment

We also looked at the behavior of the score test and of the likelihood ratio test [5] under variable weights in the empirical dataset. Table 4 displays results pertaining to the enrichment tests in the gene-set-based analyses.

**Table 4:** Results of the gene-set enrichment analysis run in the Swedish sample (N=4940; prevalence in the sample = 0.49). The 2 gene-sets included variants with MAF below 5% (A) or below 1% (B).

|  | Gene-set<br>(autosome variants in set) | weights<br>dbeta | LRT | Score |
| --- | --- | --- | --- | --- |
| A. | constrained | (1,1) | **7E-04** | 0.0031 |
|  | (63,492) | (.5,.5) | 0.1240 | 0.3444 |
|  |  | (1,25) | 0.0037 | 0.0331 |
|  | FMRP-Darnell | (1,1) | **0.0339** | 0.0577 |
|  | (72,161) | (.5,.5) | 0.1062 | 0.3384 |
|  |  | (1,25) | 0.0434 | 0.1319 |
|  | Gene-set<br>(autosome variants in set) | weights<br>dbeta | LRT | Score |
| B. | constrained | (1,1) | 0.0373 | 0.1139 |
|  | (61,269) | (.5,.5) | 0.2341 | 0.3988 |
|  |  | (1,25) | **0.0357** | 0.1293 |
|  | FMRP-Darnell | (1,1) | 0.0723 | 0.1679 |
|  | (69,668) | (.5,.5) | 0.1467 | 0.3621 |
|  |  | (1,25) | **0.0556** | 0.1668 |

From Table 4 we note that the likelihood ratio test appears more powerful than the score test across all conditions evaluated here. It is likely the combination of weight misspecification coupled with the presence of neutral variation in the target set that yielded the difference in power between the two tests. With the current sample and the likelihood ratio test with weights dbeta(1,1), the set of constrained genes showed significant enrichment for disruptive case mutations with MAF below 5% (i.e., P-value = 7e-04; see Table 4A). The score test under flat weights (i.e., dbeta(1,1)) with its associated p-value also passed the significance threshold, providing support for enrichment for disruptive rare case mutations of the constrained gene-set, although the evidence was weaker (P-value = 0.0031).

Note the difference in the strength of association of the two tests under variant weighting schemes. For instance, in the 5% MAF threshold analyses, the enrichment signal in the constrained gene-set was rendered non-significant when the dbeta(1,25) weights were used with the score test (P-value = 0.0331), and yet it reached statistical significance when the likelihood ratio test was employed instead (P-value = 0.0037). Had one relied on the score test and a default weighting scheme, the association signals in this pathway would have been missed.

The FMRP-Darnell gene-set showed no significant enrichment for rare case mutations, regardless of the test, MAF threshold and weighting schemes used. This result does not rule out the possibility that rarer variants (e.g., singletons) within the pathway play a role in the liability to schizophrenia phenotype. To implicate such variants, however, testing approaches other than those exploiting genetic similarity among the individuals are required.

The 1% MAF threshold yielded similar differences among the two tests (see table 1B). Note that the signal in the constrained gene-set no longer reached statistical significance. This result suggests that imposing this threshold probably removed from the target causal variants and so, weakened the association signal.

Summarizing, the empirical analysis showed that the choice of the test and of the weighting scheme is no trivial matter. The LRT always yielded smaller p-values than the score test, probably due to the greater sensitivity the latter has to weighed neutral variation and to model misspecification (as we found in the simulated data). We also found that either thresholding or relying on default weights would trick one into missing association signals. We elaborate on these results in the Discussion.

## 4. Discussion

We characterized the behavior of the likelihood ratio test and of the score test under weight misspecification in association studies based on the rare variant sequence kernel test. The principal finding of this study is that the likelihood ratio test is generally more robust to weight misspecification, and more powerful than the score test in such a circumstance. Our results are of interest because weight assignment is embedded within any set-based test and the true weights of the variants within the target are generally unknown.

As we found the power to be maximal under correct model specification, we next considered the issue of optimizing weighting. In the literature, weighting is mostly informed by allele frequency; frequency is taken as indicative of the strength of the purifying selection coefficient [8]. Accordingly, rarer variants are typically being assigned larger weights/contribution to the test statistic (e.g., [1]). This relationship between effect size, frequency and selection is not always straightforward, however, because it relies on assumptions about the extent of direct selection on the phenotype in question and the demographic history of the population [13, 10, 14]. Genes under weak selection may harbor rare as well as more common variants with disruptive effects [14]. Such variants with deleterious effects, escaping selection and occurring at relatively high frequencies in the population, are plausible also under strong purifying selection, as simulation studies have demonstrated [10]. Achieving maximal power when testing such regions requires adapting the weighting scheme to match the hypothesized selection. To this end, we proposed the use of a data-driven weighting approach. Our simulation results showed that such an approach maintains the power close to that of the true (i.e., correctly specified) model. When applied to real data, this approach allowed us to capture significant enrichment signal coming from variants with MAF below 5% within the constrained pathway [37]; P-value = 7e-04), lending support to the conclusion that such a variable weighting approach is likely to boost statistical power. Such adaptive approaches were also recommended by Zuk et al. (2014) and by Price et al. (2010) as being optimal for gene-based tests. Deriving weights based on allele frequency is but one of the possible ways of prioritizing the contribution to the test statistic of the variants within the target set [1]. Alternative weighting schemes that incorporate probabilities of a variant being damaging (as estimated by annotation tools such as e.g., Polyphen-2 [43] or SIFT [44] may also be considered.

It should be emphasized that our variable weighting approach renders thresholding unnecessary. Thresholding (either based on counts or on allele frequency) has been initially used in burden tests (e.g., [45, 9,10]; see also [46] for an overview on burden tests), but it has been employed also in sequence-based variance component tests (e.g., [47, 48]) for the purpose of removing neutral variation (see e.g., [8]). Yet, in our empirical analysis this practice was counterproductive: imposing the (arbitrarily chosen) 1% MAF threshold reduced the association signal in the constrained gene-set below the significance threshold. Considering common variants along with the rare ones in sequence-based kernel association tests appears to be justified for three main reasons. First, the use of variable weighting schemes is equivalent to applying variable frequency thresholds: the weights are removing from the test or favoring the contribution to the test statistic of the variants within the target set based on their frequency. Second, only the joint signal - coming from rare and more common variants - enabled us to detect significant enrichment. And third, importantly, with the current samples, our tests are mostly powered to locate regions under relatively weak selection pressures, and such regions are expected to harbour rare as well as common variants both with functional effects. To locate genes under stronger selection pressures, larger samples (see [14]) and the inclusion of more extreme weights (i.e., weights that overlook common variants and favour the rarer ones) will probably be required.

In the empirical analysis, we chose to correct out alpha in place of using permutations to compute the p-value. The data-driven weighting approach based on permutations is prohibitively slow when the number of tested variants within the target set (or the number of genes) and the sample is large. The Bonferroni correction though easier computationally, comes at a price in terms of power: the more weighting schemes one tries, the more stringent the significance threshold correction. An optimization algorithm for an optimal search for the true weights (e.g., [49] or limiting the choice of weights based on knowledge on theorized selection on each gene [14] would decrease the burden of multiple testing, and further increase power.

The score test is currently widely used in sequence-based association studies (e.g., [50, 51,52, 53] for both its computational efficiency and power [1]. Indeed, assuming correct specification, in some circumstances the score test is the most powerful test [1, 6]. However, the results provided herein showed that the likelihood ratio test has the compelling qualities of being generally more robust and more powerful under weight misspecification. This is an important result, given that, arguably, misspecified models are likely to be the rule rather than the exception in the weighting-based approaches.

## Acknowledgements

We thank to the Swedish cohort participants whose data we analyzed in this study. This research was done while Camelia Minică visited Benjamin Neale at the Broad Institute of MIT and Harvard. Camelia Minică and Jacqueline Vink are supported by the ERC starting grant 284167 (PI-JMV). For this research visit, Camelia Minică was supported also by the Talent Grant FPPT1404 offered by the Scientific and Ethical Review Board and the Faculty Board Vrije Universiteit Amsterdam. The authors wish to acknowledge the generous use of the Broad Institute compute cluster in this work.

